# An updated proteomic analysis of *Drosophila* hemolymph after bacterial infection

**DOI:** 10.1101/2024.12.18.629076

**Authors:** Samuel Rommelaere, Fanny Schüpfer, Florence Armand, Romain Hamelin, Bruno Lemaitre

## Abstract

Using an in-depth Mass Spectrometry based proteomics approach, we provide a comprehensivecharacterization of the hemolymphatic proteome of adult flies upon bacterial infection. We detected and quantified changes in abundance of several known immune regulators and effectors, including multiple antimicrobial peptides, peptidoglycan-binding proteins and serine proteases. Comparison to previously published transcriptomic analyses reveals a partial overlap with our dataset, indicating that many proteins released into the hemolymph upon infection may not be regulated at the transcript level. Among them, we identify a set of muscle-derived proteins released into the hemolymph upon infection. Finally, our analysis reveals that infection induces major changes in the abundance of proteins associated with mitochondrial respiration. This study uncovers a large number of previously undescribed proteins potentially involved in the immune response.

## Introduction

Upon systemic bacterial infection, *Drosophila* mount an immune response that culminates in the secretion of microbicidal proteins in the hemolymph^12,3^. Transcriptomic analyses of the *Drosophila* immune response have given insight into the exquisite complexity of this process^4–8^. However, these methods do not capture post-transcriptional regulation of genes involved in the immune response. Proteomic approaches aim to fill this gap, and were instrumental in early identification of several antimicrobial peptides and other immune-induced molecules^9–15^. Mass spectrometry instrumentation has greatly improved in the past decades and allows detection and quantification of proteins with much greater sensitivity. We recently used such a method to uncover changes in the *Drosophila* hemolymphatic proteome upon colonization by a bacterial endosymbiont^16^. Here we used a similar strategy to provide in-depth characterization and quantification of changes in the hemolymphatic proteome upon systemic bacterial infection.

## Material and Methods

### Fly lines and infections

Flies were maintained at 25°C on cornmeal medium (35.28 g of cornmeal, 35.28 g of inactivated yeast, 3.72 g of agar, 36 ml of fruits juice, 2.9 ml of propionic acid and 15.9 ml of Moldex for 600 ml of medium). All experiments were done on 5 day-old isogenic *w*^1118^ Drosdel adult female flies. Systemic infections with *Pectobacterium carotovora carotovora 15* (*Ecc15*) and *Micrococcus luteus* (*M. luteus*) were performed by pricking flies in the thorax with a needle previously dipped into a concentrated bacterial pellet at OD600:200^17^. Infected flies were maintained at 29°C. Hemolymph samples were collected 2, 6 and 24 hours after infection. Isogenic *Rel^E20^* flies and *spz^rm7^* flies were infected with *Ecc15* and *M. luteus* respectively, and their hemolymph was drawn at 6 hours post infection.

### Hemolymph extraction

Hemolymph was extracted using a Nanoject II (Drummond) microinjector. About 1 μL of hemolymph was collected and frozen at -80°C. Hemolymph was then diluted 10 times in PBS containing Protease Inhibitor Cocktail 1X (Roche). 1 μl was used for protein quantification with the Pierce BCA Protein Assay Kit (Thermofisher). The remaining 9 μl were mixed with SDS (0.2% final), DTT (2.5mM final) and PMSF (10μM final). Aliquots of 15μg were used for proteomics analysis.

### Sample preparation and LC-MS/MS

Each sample was digested by Filter Aided Sample Preparation (FASP)^18^ with minor modifications. Dithiothreitol (DTT) was replaced by Tris (2-carboxyethyl) phosphine (TCEP) as reducing agent and iodoacetamide by chloracetamide as alkylating agent. A proteolytic digestion was performed using Endoproteinase Lys-C and Trypsin.

Resulting peptides were recovered by centrifugation. The devices were then rinsed with 50ul of 4% trifluoroacetic acid and centrifuged. This step was repeated twice and peptides were finally desalted on C18 StageTips^19^ and dried down by vacuum centrifugation. For LC-MS/MS analysis, peptides were resuspended and separated by reversed-phase chromatography on a Dionex Ultimate 3000 RSLC nanoUPLC system in-line connected with an Orbitrap Q Exactive HF Mass Spectrometer (Thermo Fischer Scientific). A capillary pre-column (Acclaim Pepmap C18; 3 μm-100 Å; 2cm x 75 μM ID) was used for sample trapping and cleaning. Analytical separations were performed at 250 nl/min over a 260 min biphasic gradient on a 50 cm long capillary column (Acclaim Pepmap C18; 2 μm-100 Å; 50cm x 75 μM ID). Acquisitions were performed through Top N (Top10) Data-Dependent acquisition. First MS scans were acquired over an m/z window of 300 to 1600 at a resolution of 120’000 (at 200 m/z). The most intense parent ions were selected and fragmented by High energy Collision Dissociation (HCD) with a Normalised Collision Energy (NCE) of 27% using an isolation window of 1.4 m/z. Fragmented ion scans were acquired at a resolution of 15’000 (at 200 m/z) with a Maximun Injection Time of 120 ms. and selected ions were then excluded for the following 60s.

### Data analysis

Database search was performed using MaxQuant 1.5.1.2^20^ against a concatenated database consisting of the Ensemble *Drosophila melanogaster* protein database (BDGP5.25). *Ecc15* and *M. luteus* protein databases (Complete_Proteome_Erwinia carotovora subsp. carotovora_4127_Sequences AUP000031174_ecc15 and Microccus_Luteus_2207Sequences_AUP000000738_LM_Feb2017_MLUT, respectively) were used to exclude protein groups originating from bacteria. Carbamidomethylation was set as fixed modification, whereas oxidation (M) and acetylation (Protein N-term) were considered as variable modifications. Label-free quantification was performed by MaxQuant using the standard settings. The statistical analysis was performed using Perseus version 1.5.5.0 [doi: 10.1038/nmeth.3901.] from the MaxQuant tool suite. Reverse proteins, potential contaminants and proteins only identified by sites were filtered out. Protein groups containing at least 2 valid values in at least one group were conserved for further analysis. Empty values were imputed with random numbers from a normal distribution (Width: 0.5 and Down shift: 1.9 sd). A two-sample t-test with permutation-based FDR statistics (250 permutations, FDR = 0.1, S0 = 1) was performed to determine significant differentially abundant candidates. Further graphical displays were performed using homemade programs written in R [R Core Team (2017). R: A language and environment for statistical computing. R Foundation for Statistical Computing, Vienna, Austria. URL https://www.R-project.org/]. The mass spectrometry proteomics data have been deposited to the ProteomeXchange Consortium via the PRIDE partner repository with the dataset identifier PXD058897. SignalP (http://www.cbs.dtu.dk/services/SignalP/) was used to predict the presence of a signal peptide in proteins. Unsupervised hierarchical clustering was performed on statistically significant differentially abundant protein groups (significant in at least one t test analysis) using R version 3.6.2 (2020-02-29). The GOATOOLS library (version 1.0.3) in Python was used for the Gene Ontology enrichment analysis on the individual clusters^21^. Venn diagrams were generated using the online tool Venny 2.0 (https://bioinfogp.cnb.csic.es/tools/venny/index.html) and other graphs were generated using GraphPad Prism version 10.0.0.

## Results and Discussion

We analysed proteome variation of the hemolymph of 5-day-old flies after bacterial infection. Isogenic *w Drosdel* flies were left unchallenged or systemically infected with the Gram-negative bacterium *Ecc15*^22^ or the Gram-positive bacterium *M. luteus*^17^. Hemolymph was extracted at 3 different time points (2, 6, and 24hrs) using a Nanoject and the proteome was examined by liquid chromatography-tandem mass spectrometry (LC-MS/MS). To determine the fraction of the proteome regulated by the Toll and Imd pathway, we use the same approach to monitor the proteome of *Relish* (*Rel^E20^*) and *spaetzle* (*spz^rm7^*) *Drosdel* mutant flies lacking functional Imd and Toll pathways, respectively, at 6hrs post-challenge. We were able to identify between 1395 and 1655 protein groups and quantify 1291 protein groups across samples (**Table 1**). A protein group contains proteins that cannot be unambiguously distinguished by unique identified peptides and are quantified together. For ease of interpretation, protein groups will be referred to as proteins.

Comparison of the proteome of unchallenged flies with previously published data^23^ revealed that 75% of the proteome of Hartley *et al.*^23^ was present in our dataset (**Figure 1A**). Another 817 proteins not found in this previous study were additionally quantified in our experiment.

**Figure 1:**
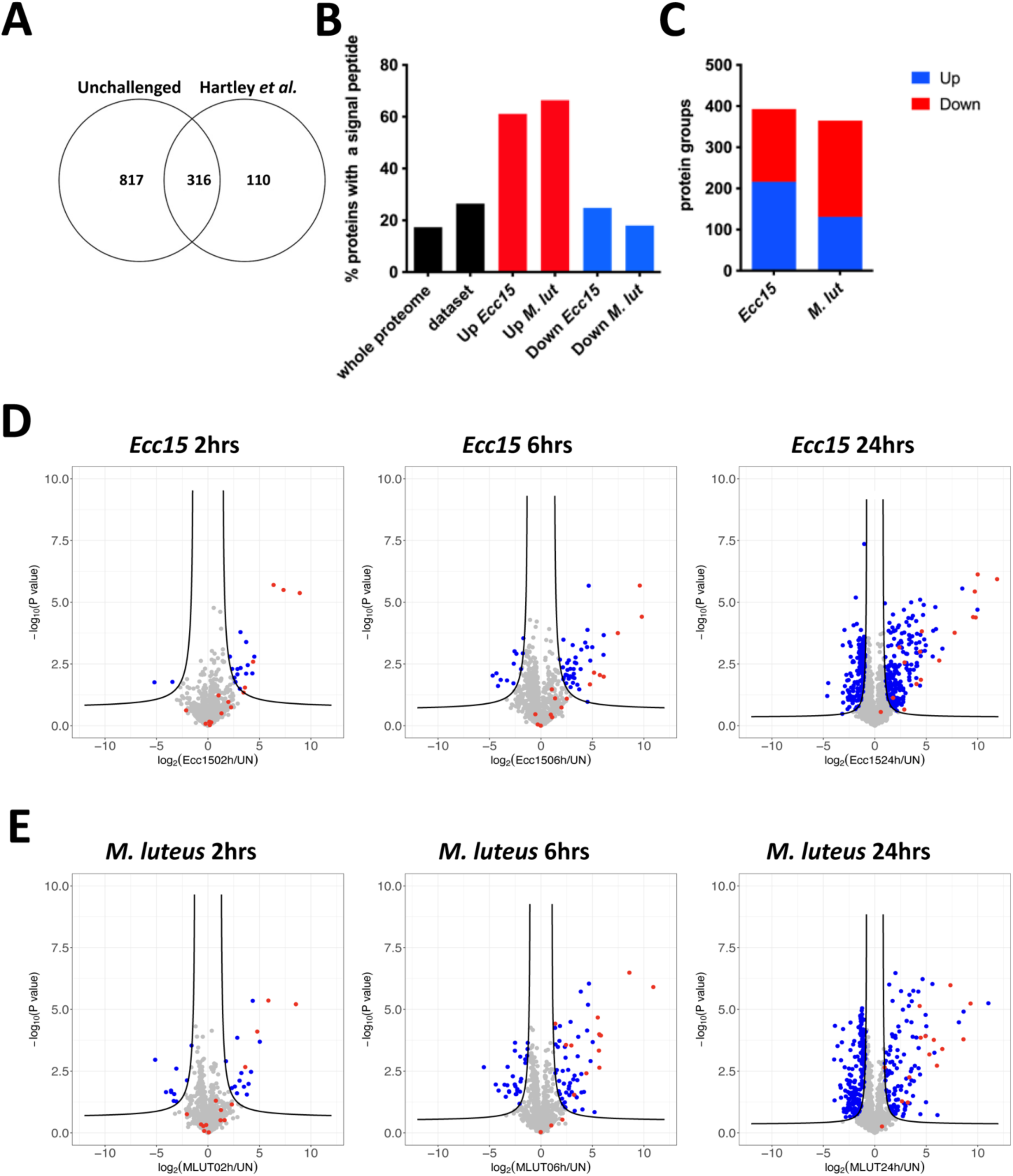
Characterization of the hemolymphatic proteome of immune-challenged flies. (A) Venn diagram showing the overlap of the proteomes described in this study (unchallenged flies) and in Hartley *et al.*^23^. (B) Proportions of proteins bearing a signal peptide as detected by SignalP algorithm. Note the enrichment in secreted proteins in the upregulated categories. (C) number of protein groups quantified and found significantly differentially expressed after infection. (D,E) Volcano plots of the differentially abundant proteins after 2 (left), 6 (middle) and 24 hours (right panel) infection with *Ecc15* (D) or *M. luteus* (E). The log_2_ fold-change indicates the differential abundance of the protein group in the infected samples versus the uninfected samples. Black lines indicate a significance cutoff, following a FDR < 0.1 and a S0 = 1. Red points highlight antimicrobial peptides.

The distribution of protein abundance was not homogenous since the 20 most abundant proteins represented 74% of the total signal. These proteins have well-established roles in lipid^24–26^ and iron transport^27^, immunity^28,29^, and energy metabolism (**Table 2**). Nine of these proteins were annotated as intracellular, suggesting that some proteins may originate from tissue or hemocyte leakage due to injury associated with the infection process or hemolymph withdrawal. Indeed, this proteome did not only contain hemolymphatic proteins; only 26.5% of the identified proteins had a predicted signal peptide and only 16.9% were associated to the Gene Ontology (GO) term ‘extracellular region’ (**Figure 1B**). These rather low numbers are regardless enriched with respect to the whole *Drosophila* proteome, as only 17.4% proteins bear a predicted signal peptide and 6.7% have the ‘extracellular’ GO term. Altogether, these data show that our method allowed robust detection and quantification of hemolymphatic proteins to an unprecedented depth.

### The immune secretome

We then interrogated the *Drosophila* hemolymphatic proteome after bacterial challenge and identified 293 and 365 differentially abundant protein groups upon *Ecc15* and *M. luteus* infection, respectively (SO=1, FDR<0.1; **Figure 1C**). Proteins with reduced abundance after infection were generally devoid of signal peptides and probably originated from tissue leakage (Figure 1B). Half of these protein groups were common to *Ecc15* and *M. luteus* infections (Figure S1A). We next focused our analysis on protein groups with increased abundance after infection. Indeed, the Volcano plots show a strong shift towards increased protein abundance upon infection, which is consistent with activation of the systemic antimicrobial response (**Figure 1D-E**). Septic injury with *Ecc15*, a Gram-negative bacterium that strongly activates the Imd pathway response, triggered upregulation of 216 proteins, while *M. luteus* infection, a Gram-positive bacterium triggering the Toll pathway, induced 131 proteins in the hemolymph compared to unchallenged conditions. In both conditions, more than 60% of the upregulated proteins had a signal peptide (**Figure 1B**) suggesting that most of the proteins induced upon infection are secreted through the canonical secretory pathway.

Several of these induced proteins were encoded by genes known to be upregulated upon systemic infection^4–6^. These proteins belong to the major humoral immune modules of *Drosophila*. All the quantified protein groups and their changes in abundance upon infection are listed in **Table 1**. We provide a non-exhaustive list of these proteins and their changes in abundance after infection.

#### Pattern recognition receptors

Several secreted Peptidoglycan Recognition Proteins (PGRPs) were found in the hemolymph (PGRP-LB, -SA, -SB1, -SC2 and -SD) and all of them were induced upon bacterial challenge in *w^iso^* but not in *Rel*-deficient Rlies. In contrast, intracellular PGRP-LE and membrane-bound PGRP-LC were not detected in our dataset, suggesting that contrary to initial reports, PGRP-LE is exclusively intracellular. GNBP1, 2, 3 and GNBP-like 3 were also detected, but only the latter was more abundant following both infections.

#### Toll-PO(phenoloxidase) serine protease signaling

We also identified a large set of immune-induced serine proteases (**Table 4**). Many of these have already been extensively characterized in the context of the immune response and participate in Toll pathway activation and/or the melanisation cascade. All the serine proteases and serine protease inhibitors involved in this pathway were detected in the hemolymph. These include: ModSP, cSP48, Grass, Psh, Hayan, Ser7, MP1, SPE, Sp7, cSPH35, cSPH242, Spn42Dd, Necrotic, Spn28Dc and Spn27A^30–37^. The abundance of half of these serine proteases increased in the hemolymph after both infections, notably cSP48, Hayan, Ser7, SPE and Sp7; the others remained stable, and only ModSP was reduced after infection. Prophenoloxidase (PPO) protein amounts were not altered by infection. As expected, known targets of the Toll pathway were not induced in *spz^rm7^* mutant Rlies (e.g. Spn28Dc and Necrotic^15,38^).

##### Antimicrobial peptides

Our proteomic analysis identified sixteen antimicrobial peptides that were more abundant in the hemolymph upon infection (**Table 3**). AttA and AttC were among the earliest and most strongly induced proteins in both *Ecc15* and *M. luteus* samples. Most antimicrobial peptides were triggered by both infection but to different extents, depending on the inducer. As previously described^39,40^, Drosomycin and several Bomanin peptides (CG15067, CG5791 & IM23) were prominently induced upon *M. luteus* infection. As expected, DptA was more strongly induced upon *Ecc15* challenge than *M. luteus* infection, and was not increased in *Ecc15*-infected *Rel^E20^* mutant Rlies^41^. This was also the case for Cecropins and IM18. Consistent with transcriptomic data^6^, DptB protein did not seem to be coregulated with DptA, and was secreted to the hemolymph in similar amounts following both infections (Figure 2A and B). The host defense peptide Defensin, which is known to be expressed at a much lower level than other inducible AMPs^42^, and several Bomanins (BomS1, S2, S3, S4, S5, S6, BomT1 and BomT3) were not detected in our dataset.

**Figure 2:**
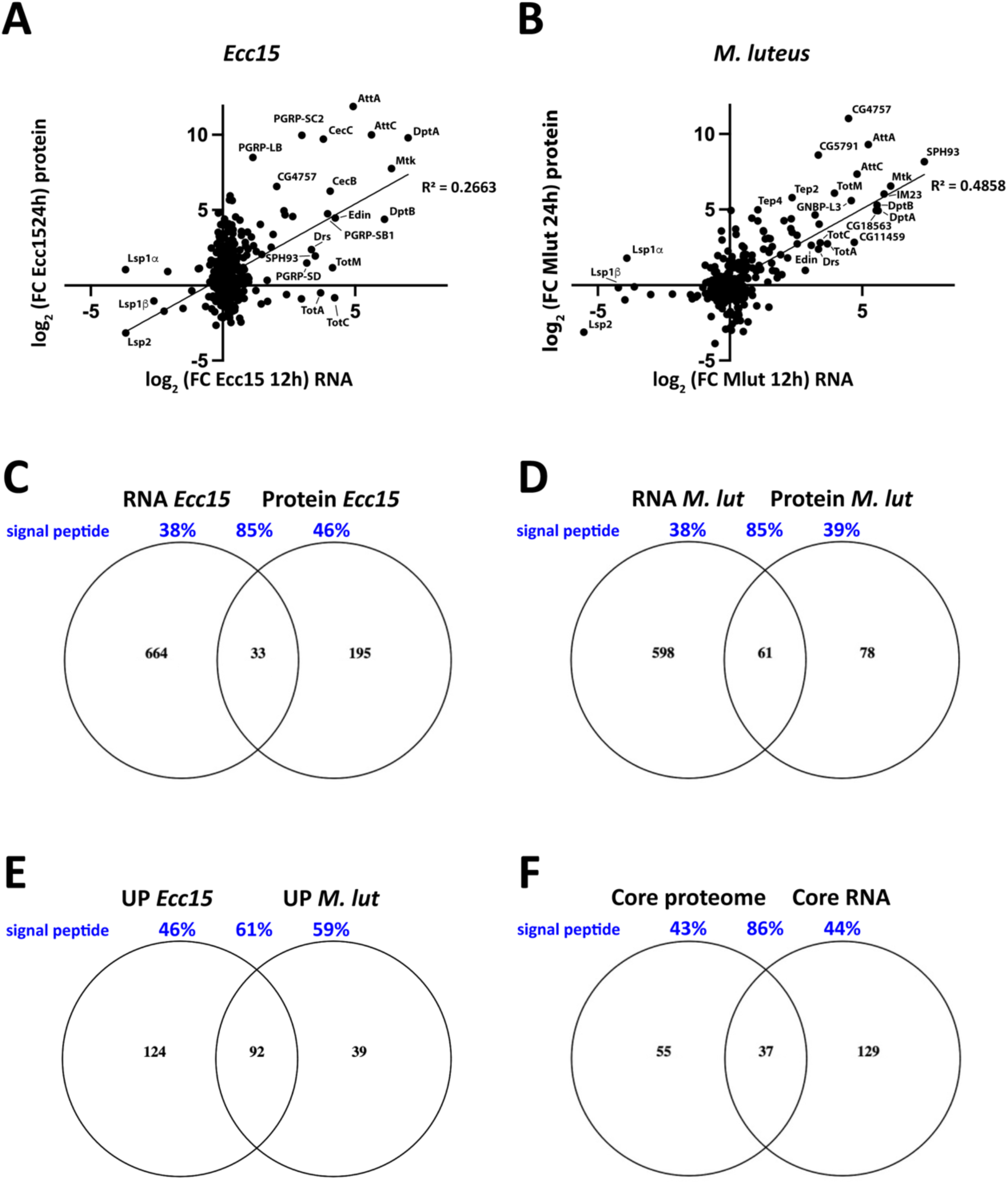
(A-B) Comparison of changes in secreted protein abundance in the hemolymph 24 hours after *Ecc15* (A) or *M. luteus* (B) infection to the change in transcript levels 12 hours after the same infection. The analysis was restricted to proteins containing a signal peptide. Data are plotted as log_2_ of the fold change (FC) of the infected over unchallenged condition. The most dysregulated genes/proteins are highlighted on the graph. (C,D) Venn diagrams showing the intersection of transcripts and proteins with increased abundance after *Ecc15* (C) or *M. luteus* (D) infection. (E) Venn diagrams indicating protein groups more abundant in the hemolymph only in response to infection with *Ecc15* (left), *M. luteus* (right) or in both infections (intersection). The latter group represents the core immune secretome. (F) Venn diagram showing the intersection of core immune genes (described in Troha *et al.*^6^) and the core immune secretome (described in panel E). Values in black represent the number of genes/proteins in each group while values in blue indicate the percentage of proteins bearing a signal peptide in the same groups.

##### Clotting factors

Clot formation prevents excessive bleeding upon injury and has microbicidal effects^15,43–45^. While most clotting studies have been done in larvae, several known components were detected in our samples of adult hemolymph, including Fondue, Hemolectin, Lsp1alpha, Lsp1beta, Lsp1gamma and Lsp2. Most of these remained stable upon infection with the exception of Lsp1a, which increased in abundance after *M. luteus* infection.

##### Iron sequestration

Iron is a key nutrient for both *Drosophila* and invading pathogens^46,47^. As a consequence, iron sequestration by the host is a strategy to prevent microbial growth. While both the iron-binding *transferrin 1* and *transferrin 3* genes are induced by Gram-positive infection, only Tsf1 protein was detected in the hemolymph and increased in abundance after infection by *M. luteus*. In contrast, the Ferritin subunits Fer1hch and Fer2lch were reduced in the hemolymph after both *Ecc15* and *M. luteus* infections.

##### Opsonins and other hemocyte binding molecules

Proteins involved in bacterial opsonisation, wasp egg encapsulation and hemocyte homeostasis were also detected. Thioester-containing proteins (Teps) play a role in cellular and humoral immunity as they promote phagocytosis of Gram-positive bacteria and Toll pathway activation^48–51^. Only Tep2 and Tep4 were detected in our dataset and both were induced after infection. Several secreted lectins were also present in the hemolymph, including Lectin-28C, Lectin-33A, Lectin-37Da and Lectin-GalC1. Finally, three secreted Nimrod B proteins were detected in the hemolymph: the adipokine NimB5 that controls hemocyte proliferation^52^ (ref) and the as-yet-uncharacterized NimB2 and NimB3 proteins.

##### Signalling molecules

We also detected several cytokines and signalling molecules. The Toll ligand Spz and the JNK ligand Eiger were present in the hemolymph; the latter was detected only after *Ecc15* infection. Although they are known to be secreted, the JAK-STAT ligands Upd1, 2 and 3 were not detected, suggesting either low expression or strong binding to tissues. Similarly, most neuropeptides were not detected, with the exception of Proctolin. Surprisingly, the Notch ligand Delta was found in the hemolymph and its abundance decreased upon both infections. These results indicate that infection by either Gram-negative or Gram-positive bacteria induced important changes in the hemolymphatic proteome. The fact that many proteins with increased abundance after infection are extracellular and have immune functions further confirms the validity of our approach.

### Comparison with the transcriptional immune response

We then compared transcriptomics data from Troha *et al.*^6^ to our proteomic dataset. We computed the fold changes in gene expression 12 hours post infection as compared to unchallenged condition and plotted it against the fold change in protein abundance 24 hours after bacterial infection (Figure 2A and B). Overall, the correlation between transcript and protein levels was weak but significant (R^2^ =0.1827 and R^2^= 0.3546 for *Ecc15* and *M. luteus* datasets, respectively, **Figure S1B and C**). There was a stronger correlation between RNA and protein levels when this analysis was carried out on secreted proteins only (R^2^ =0.2663 and R^2^= 0.4858 for *Ecc15* and *M. luteus* datasets, respectively; **Figure 2A and B**). As previously reported, these results suggest that there is no linear correlation between transcripts and protein levels^16,53^. However, the differences in sampling time (12 hours for RNA, 24 hours for proteins) or different kinetics in transcription and translation also likely contribute to these discrepancies. We then tested whether genes with dysregulated mRNA (as reported by Troha *et al.*^6^, log2 fold change <-2 or >2) were also differentially abundant at the protein level. There was no significant overlap between downregulated genes and proteins with reduced abundance (**Figure S1D**). In contrast, 14.5% of the proteins more abundant in the hemolymph after *Ecc15* infection also had increased gene expression at the transcript level, while 44.5% of the proteins induced by *M. luteus* infection were also upregulated at the mRNA level (**Figure 2C and D**). As expected, a majority of these proteins bore a signal peptide and were predicted to be secreted (85% in both *Ecc15* and *M. luteus* infection). Most of these proteins are known immune molecules (e.g., AMPs, serpins, PGRPs) and have been described above. Surprisingly, only a fraction of proteins upregulated in the proteomics dataset but not induced at the RNA level had a signal peptide (46.2% and 38.5% for *Ecc15* and *M luteus*, respectively). These observations suggest that a large proportion of proteins more abundant after infection are not regulated at the transcriptional but rather at the translational or post-translational level. Changes in abundance of signal peptide-containing hemolymphatic proteins could be caused by a regulated release through the secretory pathway while proteins without a signal peptide may be non-specifically released upon cell death and damage (see below).

### A common immune inducible secretome

We then focused our analysis on the immune secretomes that are modulated upon *Ecc15* and *M. luteus* infection. Among the induced proteins, 124 were specific to *Ecc15* infection, while 39 were induced only after *M. luteus* infection. 92 proteins were induced by both infections. We define these as the common immune proteome of *Drosophila* (Figure 2E and **table 5**). 76% of these proteins bore a signal peptide or were annotated as ‘extracellular’. This list contains 13 AMPs, 13 serine proteases, 4 serpins and 5 secreted PGRPs. Over 50% of these proteins are encoded by known immune-regulated genes or have already been linked to immunity^4–6,54^. Additionally, there was a large overlap between this common immune secretome and the common set of immune genes described by Troha et al.^6^ in their transcriptomic analysis (Figure 2F). Most of the proteins common to both groups carried a signal peptide (86.5%). However, many proteins identified in the common secretome were not reported as transcriptionally regulated and have no known role in the immune response (e.g., CG3868, QC and CG31997). In this group, 56.4% of the proteins did not encode a signal peptide and were expected to be intracellular. This included muscle-derived proteins (see below), a regulator of chromatin condensation (Df31) and proteins involved in translation, secretion and proteostasis (e.g. mRpL22, RpL26, Sar1, Ufm1). The presence of these intracellular proteins in the hemolymph is likely a consequence of infection-induced tissue damage.

### Enrichment in hemolymphatic muscle-derived proteins after infection

We then used an unsupervised hierarchical clustering strategy to classify proteins having a different relative abundance after infection with *Ecc15* (**Figure 3A**) or *M. luteus* (**Figure 3B**). This analysis revealed protein subsets with different kinetics of change in relative abundance. Gene Ontology analysis of these clusters revealed that proteins increasing in abundance after infection mostly fell into categories linked to stress and immune response and protease activity (**Figure 3A,B**).

**Figure 3:**
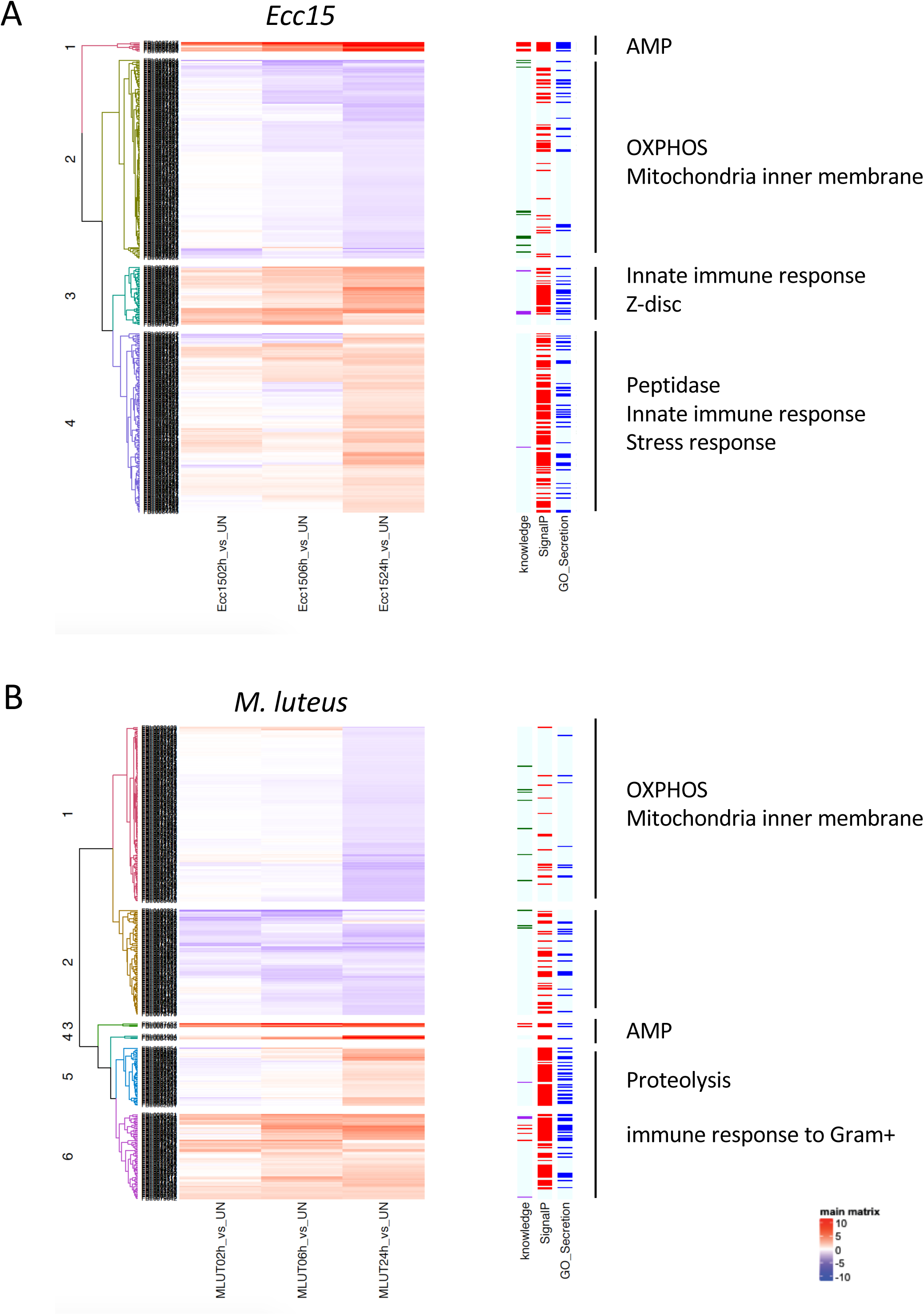
Hierarchical clustering of differentially represented hemolymphatic proteins after *Ecc15* (A) and *M. luteus* (B) infection. Left columns highlight proteins predicted to be secreted (SignalP algorithm, red bars) or annotated as secreted (GO terms: extracellular or secreted, blue bars). Representative significant enrichments for Gene Ontology terms are shown for each cluster.

Importantly, a significant enrichment in proteins associated with muscle and myofibril structure was also detected in these clusters. These proteins are devoid of signal peptides and are thus expected to be intracellular. These include Actinin, Bent, Act79B, Act88F, Mlc2, unc89 and Zasp66 (**Table 6**). Since both Rly infection and hemolymph collection are performed in the thorax, a body part mostly composed of muscles, this result may be an artefact. However, we do not favour this hypothesis. First, we found low amounts of most of these proteins in the unchallenged samples, suggesting that injury associated with hemolymph collection does not result in a massive release of muscular proteins. Second, there were increasing amounts of these proteins in the hemolymph over the course of infection. Also, muscle-borne proteins were more abundant in the hemolymph when Rlies were infected with a pathogenic bacterium (*Ecc15*) as compared to a relatively innocuous microbe (*M. luteus*). Finally, infected *spz^rm7^* and *Rel^E20^* mutant Rlies had more muscle-derived proteins in their hemolymph than their control counterparts at 6 hours post-infection, suggesting that uncontrolled bacterial growth leads to leakage of muscle proteins. Consistent with our analysis, the cytoskeletal protein Actinin is thought to function as a Damage Associated Molecular Pattern that activates the JAK-STAT ligand upd3, a cytokine that orchestrates the systemic wound response^55–57^. Based on these results we hypothesize that muscle proteins are released in the hemolymph as a consequence of the injury and escalated by bacteria-induced cytolysis. It would be interesting to test if these other muscle proteins could also act as DAMPs in this context.

### Infection decreases hemolymphatic mitochondrial proteins

Several mitochondrial proteins were detected in the hemolymph in unchallenged conditions. Such proteins are not expected to be secreted, although this finding has already been reported^58,59^. Because many muscle-derived proteins were also detected in our dataset, we hypothesized that these mitochondrial proteins originate from muscle tissue and were released during the extraction procedure. Surprisingly, and contrary to the pattern observed for sarcomeric proteins, the mitochondrial proteins were significantly less abundant in the hemolymph after infection. Indeed, the GO analysis revealed that many protein groups whose relative abundance was reduced after infection were strongly linked to the oxidative phosphorylation process and the mitochondrial inner membrane (GO analysis, *p*<10e-40). Indeed, 33% and 26% of the downregulated protein groups were of mitochondrial origin in both *Ecc15* and *M. luteus* samples, respectively. Although Cytochrome c was one of the earliest downregulated proteins after both bacterial challenges, most mitochondrial proteins became significantly less abundant 24hrs post-challenge, suggesting important changes in mitochondrial function, probably in the muscles. This reduction could be associated with muscle damage and may be a direct consequence of the injury, the bacterial infection or the immune response itself. Indeed, mitochondrial membranes are highly enriched in the negatively charged phospholipid cardiolipin and it has been suggested that cationic AMPs might disrupt these organelles^60,61^. However, we do not favor this hypothesis as *Rel^E^*^20^ mutants lacking a large subset of AMPs display an even deeper depletion of mitochondrial proteins after infection, suggesting that uncontrolled bacterial proliferation rather than AMP-mediated damage could be a cause of mitochondrial protein reduction. Consistent with this idea, mitochondrial protein depletion is more pronounced upon infection with the pathogenic bacteria *Ecc15* than after infection with the innocuous microbe *M. luteus*. Alternatively, reactive oxygen species produced during the immune response could alter respiratory complexes or damage mitochondria. Further work is needed to delineate the impact of AMPs, ROS and infection on mitochondrial membranes in the context of infection.

### Concluding remarks

Altogether, our dataset (Table 1 and ProteomeXchange dataset identifier PXD058897) provides an extensive characterization of the hemolymphatic proteome and its changes upon systemic bacterial infection. This study confirms the induction of previously described immune-induced proteins and identifies interesting new candidates that were not detected in transcriptomic analyses, reinforcing the idea that proteomics represents a valid and complementary approach to study the immune response.

## Data Availability Statement

The data that support the findings of this study are openly available in ProteomeXchange repository with the dataset identifier PXD058897

## Acknowledgements

We thank Florent Masson for fruitful discussions and Hannah Westlake for critical review and editing of the manuscript. This study was funded by SNFS grants 310030_215073 and CRSII5_186397

---

**Figure S1:**
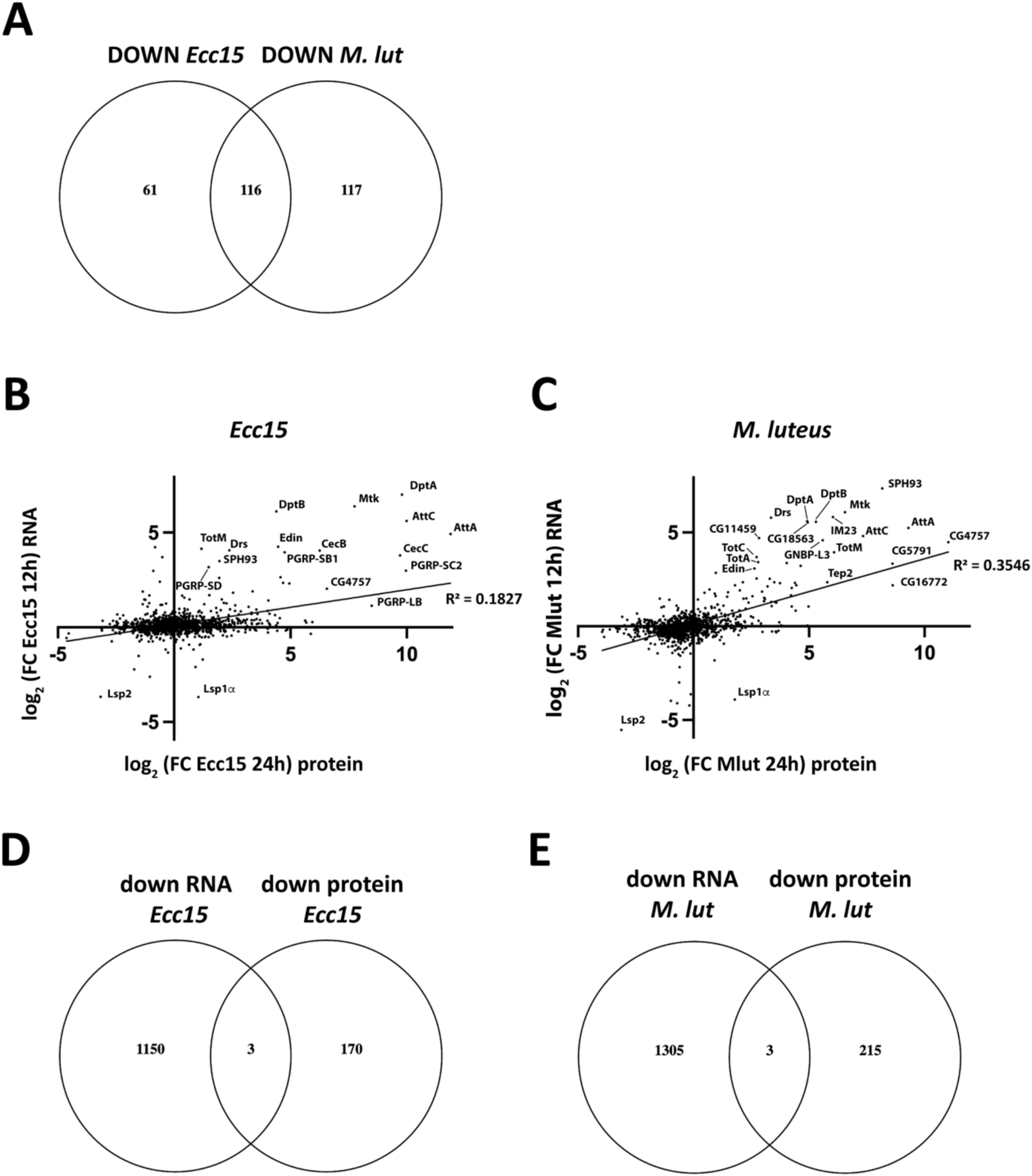
(A) Venn diagram showing the overlap of protein groups with decreased abundance after infection with *Ecc15* (left) or *M. luteus* (right). (B,C) Comparison of changes in protein abundance in the hemolymph 24 hours after *Ecc15* (B) or *M. luteus* (C) infection to the change in transcript levels 12 hours after the same infection. Data are plotted as log2 of the fold change (FC) of the infected over unchallenged condition. The most highly dysregulated genes/proteins are highlighted on the graph.

